# Direct retino-iridal projections and intrinsic iris contraction mediate the pupillary light reflex in early vertebrates

**DOI:** 10.1101/2024.02.29.582767

**Authors:** Cecilia Jiménez-López, Paula Rivas-Ramírez, Marta Barandela, Carmen Núñez-González, Manuel Megías, Juan Pérez-Fernández

## Abstract

The pupillary light reflex (PLR) adapts the amount of light reaching the retina, protecting it and improving image formation. Two PLR mechanisms have been described in vertebrates. First, the pretectum receives retinal inputs and projects to the Edinger-Westphal nucleus (EWN), which targets the ciliary ganglion through the oculomotor nerve (nIII). Postganglionic fibers enter the eye-globe, travelling to the iris sphincter muscle. Additionally, some vertebrates exhibit an iris-intrinsic PLR mechanism mediated by sphincter muscle cells that express melanopsin inducing muscle contraction. Given the high degree of conservation of the lamprey visual system, we investigated the mechanisms underlying the PLR to shed light onto their evolutionary origins. Recently, a PLR mediated by melanopsin was demonstrated in lampreys, suggested to be brain mediated. Remarkably, we found that PLR is instead mediated by direct retino-iridal cholinergic projections, a mechanism not demonstrated before, although suggested to be present in mice. This retina-mediated PLR acts synergistically with the iris-intrinsic mechanism mediated by melanopsin, which has contribution of gap junctions, as in other vertebrates. In contrast, we show that lampreys lack the brain-mediated PLR. Our results suggest that two eye-intrinsic PLR mechanisms were present in early vertebrate evolution, whereas the brain-mediated PLR has a more recent origin.

## Introduction

Pupil contraction through the pupillary light reflex (PLR) reduces the amount of light that reaches the retina, protecting it and maximizing image formation efficiency^1–2^. In vertebrates, two different PLR mechanisms have been described. In the first one, the retina sends light information through the optic nerve (nII) to the pretectum, a brain area that, in turn, projects to the Edinger-Westphal nucleus (EWN). Then, the EWN sends signals to the ciliary ganglion through the oculomotor nerve (nIII), and efferent fibers from the ciliary ganglion enter the ocular globe, travelling to the iris sphincter muscle and releasing acetylcholine (Ach)^1^. Additionally, fish, amphibians, birds, and nocturnal/crepuscular mammals exhibit a PLR mechanism that is intrinsic to the iris, mediated by sphincter muscle cells that express melanopsin and can thus act as photoreceptors and evoke muscle contracion^3–6^. These melanopsin-expressing muscle fibers are in turn connected to adjacent muscle fibers via gap-junctions, allowing the spread of the excitability changes evoked by light^5^. In mice, it has been shown that a cholinergic component can still contribute to the PLR in the isolated eye^7^. Thus, it has been suggested that light information from the retina can target the iris without entering the brain, possibly through the ciliary ganglion or via direct projections from the retina to the iris^7–9^.

However, this mechanism is controversial and the precise pathway has not been determined yet ^5^. Moreover, the evolutionary origin of the PLR mechanisms is also unknown. Recently, it has been demonstrated that lampreys, belonging to the oldest group of extant vertebrates, also exhibit PLR that is mediated by melanopsin^10^. Preliminary experiments suggested that the lamprey PLR is brain-mediated since it could not be evoked in isolated eyes^10^. Lampreys have a well-developed visual system with image-forming camera eyes, and an organization of eye muscles remarkably similar to that of other vertebrates. Moreover, the main visual centers in the brain are also present in these animals^11–16^. Therefore, we investigated the mechanisms underlying the PLR to shed light onto their origin and evolution in vertebrates. Remarkably, we found that the PLR is mediated by direct cholinergic projections from the retina to the iris. Additionally, a second mechanism, intrinsic to the iris, also contributes synergistically to the PLR. This iris-intrinsic mechanism is mediated by melanopsin, and gap junctions contribute to the spread of the excitability, therefore being like the mechanism reported in other vertebrates^5^. However, we here show that the brain-mediated PLR is not present in lampreys. Our results therefore suggest that two PLR mechanisms intrinsic to the eye were present at the dawn of vertebrate evolution and were conserved in some vertebrate groups, whereas the brain-mediated PLR is evolutionarily more recent.

## Results

### Characterization of the lamprey iris sphincter muscle

The presence of a PLR in lampreys^10,17^ implied that intraocular muscles would also exist, but their presence had only been suggested in the species *Mordacia mordax*^18^. Sagittal sections of the eye showed that the iris sphincter muscle is formed by a thin 2-3 layer of fusiform cells adjacent to the external side of the pigmented layer of the iris (Figs. 1a-c) in the two species analyzed, *Petromyzon marinus* and *Lampetra fluviatilis*. Transmission electron microscopy confirmed that numerous myofilament fiber bundles are present in these cells (Fig. 1d).

**Fig 1.**
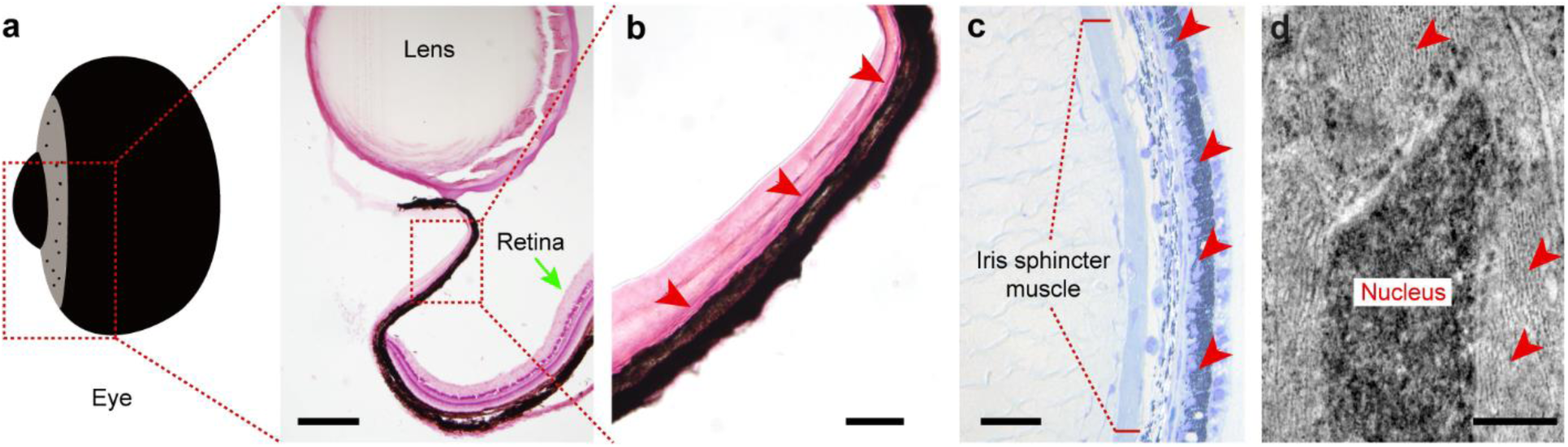
Characterization of the iris sphincter muscle. **a**, Hematoxilin-eosin stained sagittal section of the eye, showing the overall anatomy and location of the iris sphincter muscle. **b**, Detail of the photomicrograph shown in **a**, indicating the location of the sphincter muscle fibers in the iris (red arrowheads). **c**, Toluidin blue stained sagittal section of the eye showing that the iris sphincter muscle is formed by a thin layer of fibers close to the pigmented layer of the iris (red arrowheads). **d**, Transmission electron microscopy image showing the presence of abundant bundles of myosin fibers (red arrowheads). Scale bars = 250 µm in **a**; 50 µm in **b**; 100 µm in **c**; 0.2 µm in **d**.

### Lampreys lack a brain-mediated PLR

Lampreys have a highly conserved visual system^11–16^, and it has been suggested that their PLR is mediated by the brain^7^. Thus, we first investigated whether the pretectum-Edinger-Westphal nucleus pathway is also present in lampreys. We used an isolated eye/brain preparation to monitor iris contraction while manipulating/stimulating brain regions of interest^11,14,19^. In this preparation, reliable pupil contraction (tracked with DeepLabCut^20^) was evoked by presenting light with an LED to one eye (Fig. 2a). The PLR was analyzed in *P. marinus*, and *L. fluviatilis* (Figs. 2b-f) and contraction rates were like those previously reported in intact animals^10^. Pupil contraction was somewhat smaller in *L. fluviatilis* (Figs. 2e,f), but since no obvious differences were observed between the two species, the mechanisms described apply to both.

**Fig 2.**
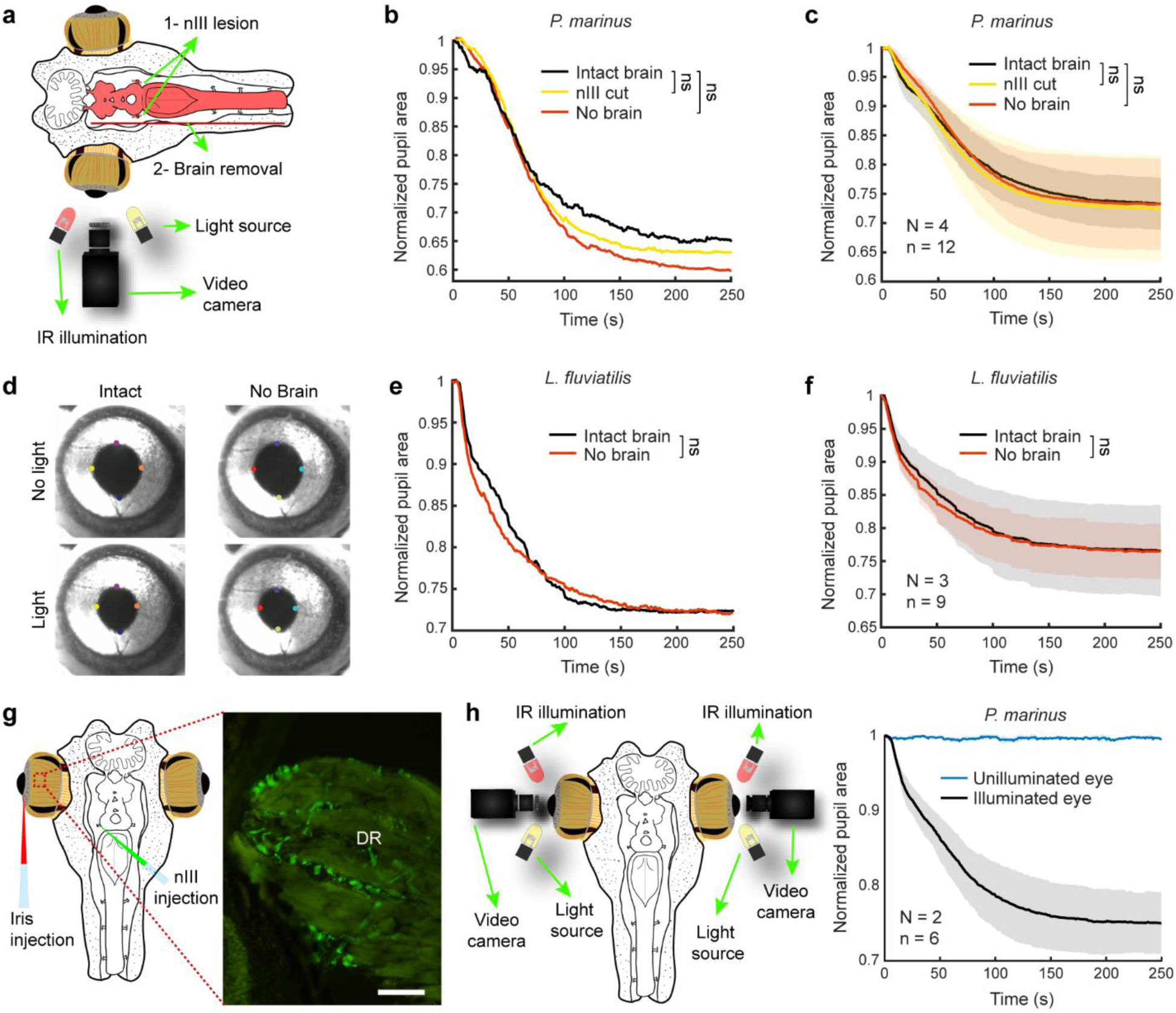
Lampreys lack brain-mediated PLR. **a**, Experimental preparation used to investigate the role of the brain in the PLR. **b**, Normalized pupil area in response to light stimulation in the intact preparation (black line), after sectioning both oculomotor nerves (nIII, yellow), and after brain removal (red), in a representative *Petromyzon marinus* specimen (paired t-test p=1). **c**, Normalized averaged data (mean ± SD) for pupil area in response to light stimulation in the intact preparation (black line), after bilateral nIII sectioning (yellow), and after brain removal (red) in *Petromyzon marinus* (N=4, paired t-test p=1, see Extended Data Fig. 1). **d**, Frames showing pupil size before (top) and after (bottom) light stimulation, in a representative animal before (left) and after brain removal (right). **e**, Normalized pupil area in response to light stimulation in the intact preparation (black line), and after brain removal (red), in a representative *Lampetra fluviatilis* specimen (paired t-test p=0.734). **f**, Normalized averaged data (mean ± SD) for pupil area in response to light stimulation in the intact preparation (black line), and after brain removal (red) in *Lampetra fluviatilis* (N=3; paired t-test p=1). **g**, Dual tracer injections performed to investigate putative connections between the iris and the brain. No retrogradely labeled ciliary ganglion cells were found (see Extended Data Fig. 2a), although abundant terminals were observed in the muscles innervated by the nIII, including the dorsal rectus (DR, right). **h**, Preparation used to test the consensual PLR in lampreys (left). Normalized averaged data (mean ± SD, right) for pupil area in response to light to one eye. The blue line shows pupil area for the unilluminated eye, and the black line for the illuminated eye (N=2). Scale bar = 100 µm in **g**. n.s., not significant.

In other vertebrates, the fibers from the EWN exit the brain through the nIII^1,21^. Thus, we first compared pupil contraction before and after sectioning the nIII bilaterally, but no significant PLR reduction was observed (Figs. 2b,c; paired t-test p=1 for both). We also performed electrical stimulation of the nIII aimed to evoke pupil contraction (N=3). However, neither long stimulation trains with short pulses (30 s stimulation, 10 ms duration pulses, 10 Hz), nor a single long pulse stimulation (one pulse of 10 s duration) resulted in changes in pupil diameter or EMG activity in the iris, despite the activation of the extraocular muscles^22^ (Extended Data Fig. 1). In other vertebrates, the nIII targets the ciliary ganglion, which in turn projects to the iris sphincter muscle^1,21^. We applied neurobiotin to the nIII to anterogradely label putative terminals innervating the ciliary ganglion, and dextran-rhodamine in the iris to retrogradely label the ciliary ganglion cells (Fig. 2g left). Sagittal sections of the eyes together with their surrounding tissues were analyzed but no ganglionic neurons could be found, although clear anterogradely labeled terminals could be observed targeting the muscles innervated by the nIII^22^ (Fig. 2g right). However, no terminals were found in other areas. Additionally, sagittal sections were made of the intact head, following the nIII from its exit in the brain to the eye, but no evidence of a ciliary ganglion was found, fully confirming its absence in lampreys (N=2; Extended Data Fig. 2a). We also investigated the presence of a consensual PLR (N=2; Fig. 2h left). However, only the illuminated eye underwent pupil constriction (Fig. 2h right), as previously reported^10^. These results indicate that the EWN-mediated mechanism known in other vertebrates is not present in lampreys, and that lampreys lack a ciliary ganglion as previously suggested^23^. No reduced pupil contraction was observed after total removal of the central nervous system to isolate the eye (N=7; Figs. 2b-f; paired t-test p=1 in b, c, f and p=0.734 in e), fully confirming that the PLR observed in lampreys is intrinsic to the eye.

### The isolated iris shows PLR mediated by melanopsin

In other vertebrates, a mechanism intrinsic to the iris has been reported, in which the muscle fibers express melanopsin and act as photoreceptors contracting the pupil directly proportional to the amount of light^3–5^. We investigated this in lampreys, isolating the iris and applying light (N=37; Fig. 3a), which resulted in a clear PLR (Fig. 3b), suggesting that, as in other vertebrates, muscle fibers express melanopsin and mediate pupil reduction. Since it is difficult to isolate the fragile iris without affecting its integrity, small parts of the retina remained in some cases. Thus, we isolated the iris in the presence of atropine (1 mM), an antagonist of muscarinic ACh receptors, to block cholinergic transmission and prevent any possible influence of the retina (see below; Figs. 3c-f). Reduction rates were in general larger than those observed in intact eyes, most likely because the iris sphincter muscle must overcome less resistance and can thus achieve a larger contraction. The iris-intrinsic PLR is mediated by melanopsin in other vertebrates^4,6,24^, which has been shown to be expressed in the fibers of the iris sphincter muscle.^5^ Although melanopsin expression was previously reported in the lamprey retina^25^, its presence in the iris was unknown. However, the presence of an iris-intrinsic PLR in lampreys, and previous experiments showing that the lamprey PLR achieves its maximum contraction within the absorption wavelength range of melanopsin^10^, suggest that melanopsin also mediates the lamprey iris-intrinsic PLR. Thus, we analyzed the PLR in the isolated iris after blocking melanopsin with AA92593 (60 µM), a selective and reversible antagonist of melanopsin-mediated phototransduction (N=10). Light stimulation resulted in clear contraction (Fig. 3d, black line) that was significantly reduced after the application of the melanopsin antagonist (Fig. 3d, magenta line; paired t-test p<0.001). These effects were reverted after washout of the melanopsin antagonist (Fig. 3d, green line). These results indicate that, as in mice^5^, melanopsin is expressed in the muscle fibers of the iris sphincter muscle and mediates a PLR. Our results show that this mechanism was already present before lamprey divergence and was maintained in several vertebrate groups whereas in others, including humans, has been lost^4,26^.

**Fig 3.**
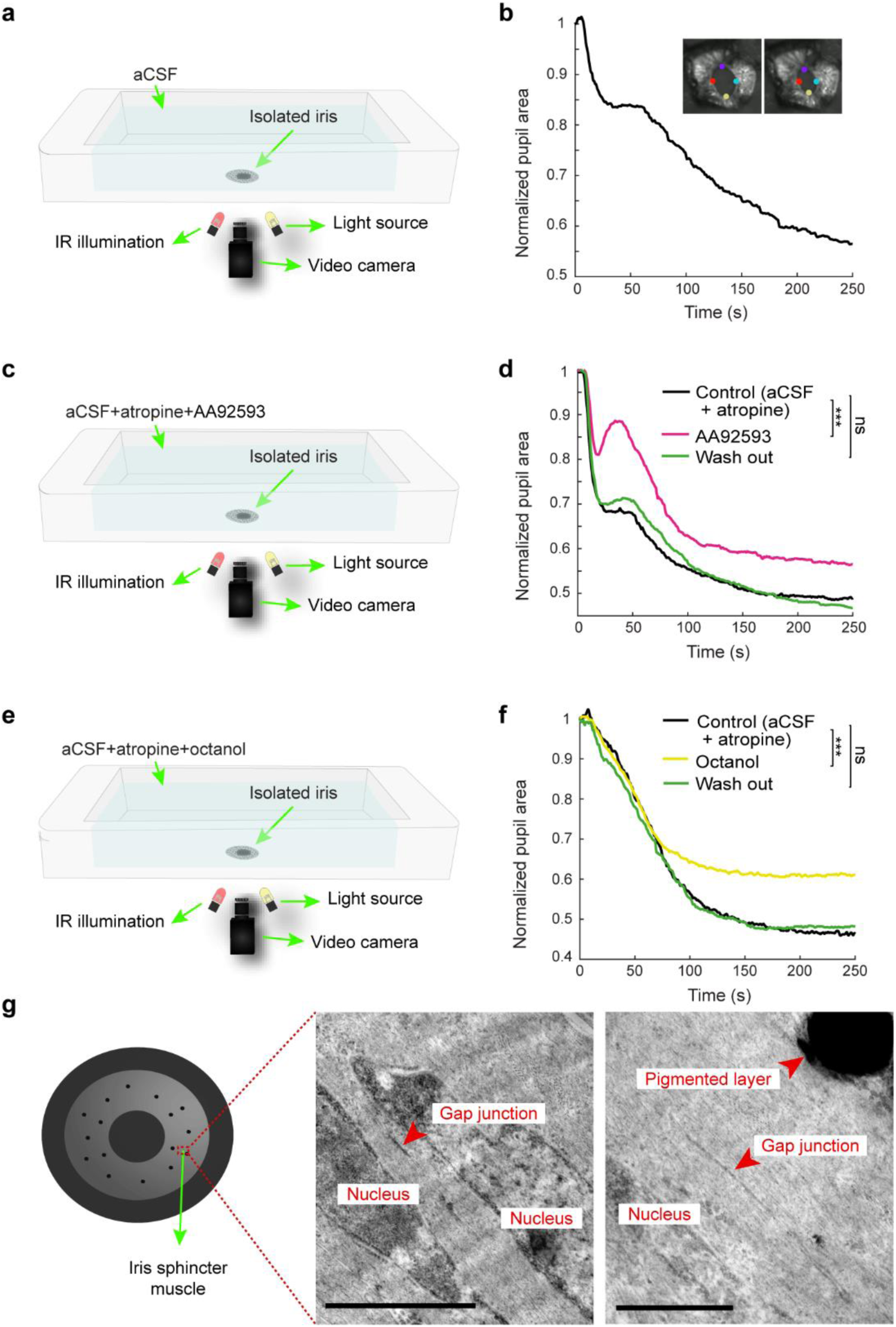
Lampreys possess a melanopsin-dependent iris-intrinsic PLR. **a**, Experimental setup to analyze pupil contraction in the isolated iris, submerged in artificial cerebrospinal fluid (aCSF). **b**, Graph showing a representative normalized pupil area through time in response to light stimulation to the isolated iris. **c**, Experimental setup to evaluate the role of melanopsin in the isolated iris. aCSF with atropine to block any potential contribution of the retinal cholinergic pathway was used as a control, and AA92593 was used to block melanopsin. **d**, Graph showing the normalized pupil area through time in response to light stimulation to the isolated iris in control conditions (black line), after applying the melanopsin antagonist AA92593 (magenta line), and after washout (green line; paired t-test *** p<0.001). **e**, Experimental setup to pharmacologically test the contribution of gap junctions to the PLR in the isolated iris. aCSF with atropine to block any potential contribution of the retinal cholinergic pathway was used as a control, and octanol was used to block gap junctions. **f**, Graph showing the normalized pupil area through time in response to light stimulation to the isolated iris in control conditions (black line), after applying the gap junction blocker octanol (yellow line, paired t-test *** p<0.001), and after washout (green line). **g**, Transmission electron microscopy images showing the presence of gap junctions between adjacent muscle fibers of the iris sphincter muscle. Scale bars = 1 µm in **g** left and 0.5 µm in **g** right.

### Gap junctions contribute to the contraction of the iris sphincter muscle

In mice, gap junctions between adjacent muscle fibers in the iris transfer changes in excitability in response to light to other muscle fibers^5^. To test whether gap junctions play a role in the lamprey PLR, we blocked them pharmacologically in the isolated iris using octanol (N=3; Fig. 3e; 0.5 mM) as previously reported in mice^5^. Again, atropine was used to block any possible contribution of the ACh retinal mechanism described below. As shown in the representative example of Fig. 3f, pupil contraction (black line) was significantly reduced after blocking gap junctions (yellow line; paired t-test p<0.001), indicating that these intercellular channels participate in the transmission of the excitability changes triggered by melanopsin. The effects exerted by the gap junction blocker were reverted after washout (Fig. 3f, green line). The presence of gap junctions in the fibers of the iris sphincter muscle was confirmed by using transmission electron microscopy (N=3; Fig. 3g). These results show that the participation of gap junctions in the iris sphincter muscle contraction was already present in early vertebrates, and further reinforce that the iris-intrinsic PLR mechanism described in mice was already present at the base of vertebrate evolution.

### Retinal cells provide cholinergic innervation of the iris

In mice, it has been suggested that intrinsically photosensitive retinal ganglion cells (ipRGCs) can evoke pupil contraction via direct cholinergic projections to the iris^2,7–9^. Although this mechanism has only been suggested in mice, we decided to test this possibility in the lamprey. We first isolated the iris and applied ACh (100 µM; N=9), that resulted in a clear pupil contraction (Fig. 4a). Given the absence of a ciliary ganglion and any brain-mediated PLR (see above), these results indicate that there may be an ACh release in the iris, likely via axons originating within the eye. To confirm the presence of an eye-intrinsic mechanism mediated by ACh, we isolated the eye and analyzed the PLR before and after the application of atropine (N=5; Fig. 4b; 1 mM). As shown in Fig. 4b (left), the PLR (black line) was significantly reduced (paired t-test p<0.001) when cholinergic transmission was blocked (orange line). These results suggested that an axonal component innervating the iris sphincter muscle is present in lampreys. To investigate the presence of axons in the iris, we immunostained for acetylated tubulin, and observed axons coursing from the retinal region to the iris both in the flat-mounted eye (Fig. 4c and Extended Data Fig. 2b) and in sagittal sections (Fig. 4d). We next aimed to uncover the origin of these projections. For this, we first applied neurobiotin in the iris and found retrogradely labeled neurons in the retina (N=11). The lateral distribution of labeled cells throughout the retina was homogeneous, and no obvious regionalization of labeled cells was observed. Neurobiotin labeled cells were found isolated (Extended Data Fig. 2c), but in some cases clusters of numerous retrogradely labeled cells were observed, both in close apposition (Fig. 4e) and forming groups of more separated cells (Fig. 4g). The low molecular weight of neurobiotin (287 Da), allows this tracer to cross gap junctions^27^, which are known to be present in the lamprey retina^28^. Thus, we hypothesized that some of the labeled cells were not retrogradely labeled, but rather electrically coupled to cells projecting to the iris. To confirm this, we did injections both combining neurobiotin and dextran-rhodamine (N=2; Fig. 4f), and only with dextran-rhodamine (3000 Da; N=6; Fig. 4h,i). The high molecular weight of dextran-rhodamine does not allow this molecule to cross gap junctions and, accordingly, no dextran-rhodamine retrogradely labeled cell clusters as those observed with neurobiotin were found, thus indicating that cells projecting to the iris are electrically coupled to other cells in the retina. As for neurobiotin application, no obvious lateral regionalization was found, and iris projecting cells (retrogradely labeled with dextran-rhodamine) were homogeneously distributed throughout the retina. In Fig. 4f, a representative example is shown of a retrogradely labeled cell, both with neurobiotin and dextran-rhodamine. Iris-projecting neurons were more abundant in the inner nuclear layer (INL) region proximal to the inner plexiform layer (IPL; Figs. 4f,i; see 29-32 for lamprey retinal organization). In the INL, iris-projecting cells were also found in the layer of horizontal cells (Fig. 4i, blue arrowhead), and, occasionally, iris-projecting cells were also present in the IPL (not shown). Cells projecting from the retina were always found in the IPL and INL. The same applied to those cells labeled with neurobiotin (including both iris projecting and electrically coupled cells), indicating that the cells electrically coupled with those projecting to the iris are also located in these two layers. Interestingly, the clusters of neurobiotin labeled cells in close apposition were always found in the INL, in the layer of inner horizontal cells (Fig. 4e). Remarkably, melanopsin expression in the lamprey retina was reported both in the INL and the IPL^25,31^, and cholinergic neurons in lampreys were also reported in these two layers^33^. Nearly all cells in the INL layer of horizontal cells express melanopsin^25,31^, indicating that both the cell clusters of electrically coupled cells and those projecting to the retina express melanopsin in this region. Additionally, melanopsin expression was also reported in the INL proximal region to the IPL, and in the IPL itself^25,31^. The presence of melanopsin in the same regions where iris-projecting cells are located suggests that iris-projecting cells likely express melanopsin, although additional experiments are necessary to confirm this. Regarding ACh, choline acetyltransferase immunoreactivity was shown in both the IPL and the INL^33^. In the INL, most cholinergic cells were reported in the layer of cells proximal to the IPL, in the same location where iris-projecting cells are more abundant, in agreement with our pharmacological results showing that iris-projecting cells are cholinergic. Altogether, these results indicate that lampreys have direct cholinergic projections from the retina to the iris that can evoke pupil contraction. Considering previous results^25,31^, cholinergic cells also express melanopsin and are in turn electrically coupled via gap junctions to other melanopsin-expressing cells.

**Fig 4.**
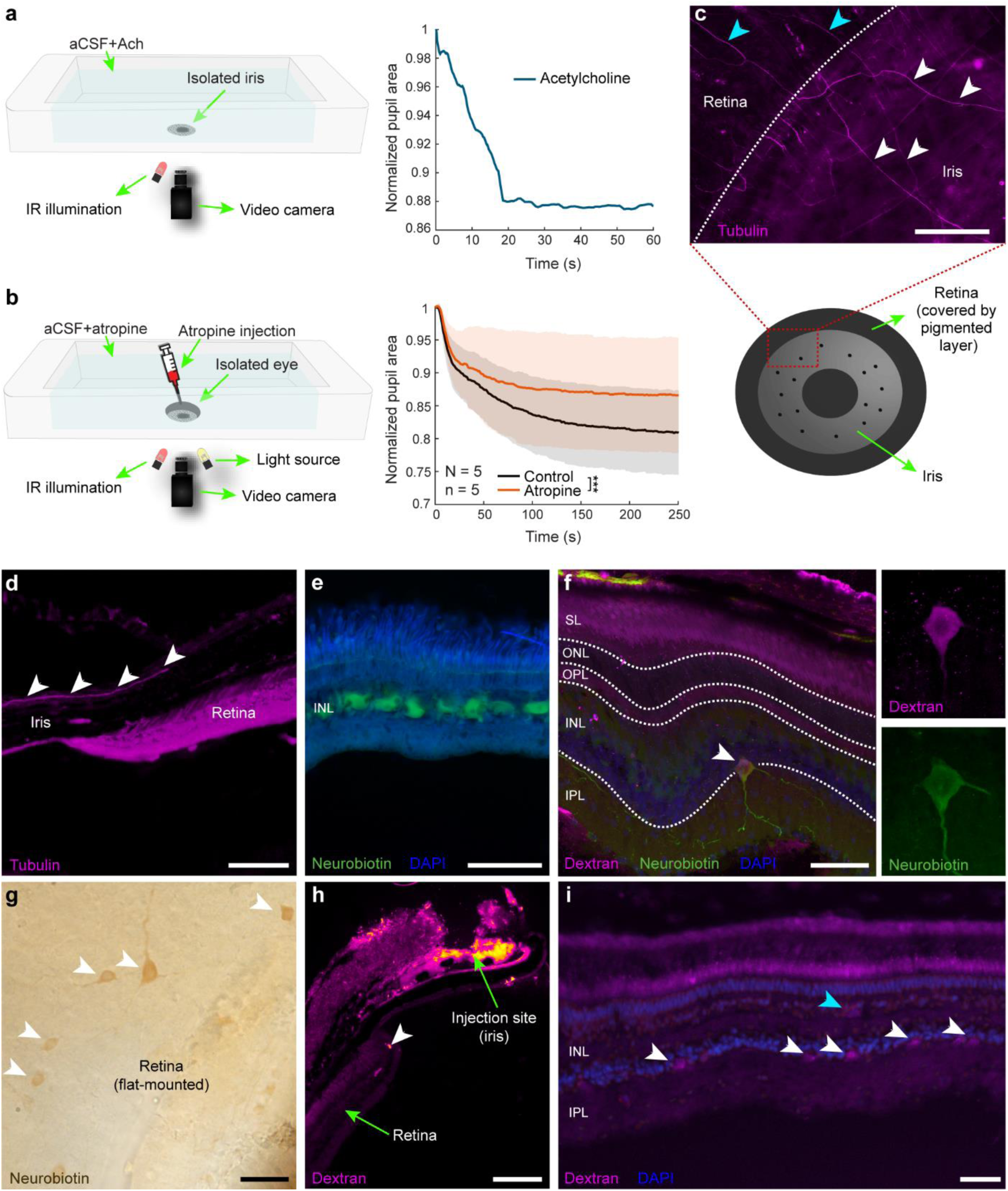
A direct projection from the retina to the iris contributes to PLR. **a**, Experimental setup to analyze pupil contraction in the isolated iris, submerged in aCSF (left), and a representative normalized pupil area in response to acetylcholine (ACh) application (right). **b**, Experimental setup (left) to analyze the contribution of ACh to the PLR in the isolated eye, using the cholinergic antagonist atropine, both bath-applied and injected in the eye. Normalized averaged data (mean ± SD, right) for pupil area in response to light stimulation in control conditions (black line), and after atropine application (orange line). ***p < 0.001 (paired t-test). **c**, Photomicrograph showing the expression of acetylated tubulin in a flat-mounted eye. Labeled axons are coursing from the retina (blue arrowheads) to the iris (white arrowheads). The location of the image can be seen in the schematic below. See also Extended Data Fig. 2b. **d**, Sagittal section of the eye showing acetylated tubulin labeling in the retina, and axons towards the iris (white arrowheads). **e**, Neurobiotin labeled cluster of neurons located in the INL, in the layer of horizontal cells. See also Extended Data Fig. 2c. **f**, Cell labeled both with neurobiotin and dextran-rhodamine. In the left, the labeling for each of the tracers is shown. **g**, Neurobiotin labeled cells in a flat-mounted retina. **h**, Representative injection site of dextran-rhodamine in the iris and retrogradely labeled cell can be seen in the retina (white arrowhead). **i**, Dextran-rhodamine labeled cells in the inner nuclear layer (INL) of the iris, both proximal to the inner plexiform layer (IPL; white arrowheads) and in the layer of horizontal cells (blue arrowhead). Scale bars = 100 µm in **c**, **e** and **h**; 50 µm in **d** and **f**;.20 µm in **g** and **i**. SL, segment layer; ONL, outer nuclear layer; OPL, outer plexiform layer.

### The iridal and retinal PLR mechanisms complement each other

The analysis of the PLR at a 15 Hz framerate allowed us to uncover the details of pupil contraction dynamics. We noticed that, in general, pupil size reduction was not constant, but in many eye-intact experiments we noticed a clear decrease in the reduction rate occurred after a few seconds. To confirm this decrease of pupil reduction, we analyzed the presence of significant slope changes (see Methods). In the graph of Fig. 5a the significant slope changes are shown for the averaged data of 7 intact eyes. At ∼ 17.6 s, the speed of pupil reduction decreased, and around 33.2 s increased again. Interestingly, in this time window, pupil contraction not only slowed down but reverted giving rise to pupil dilation when light stimulation was presented to the isolated iris (Fig. 5b). This shows that the iris-intrinsic mechanism alone has some limitations to evoke a continuous pupil reduction, but this limitation is counteracted by the cholinergic projections from the retina.

**Fig 5.**
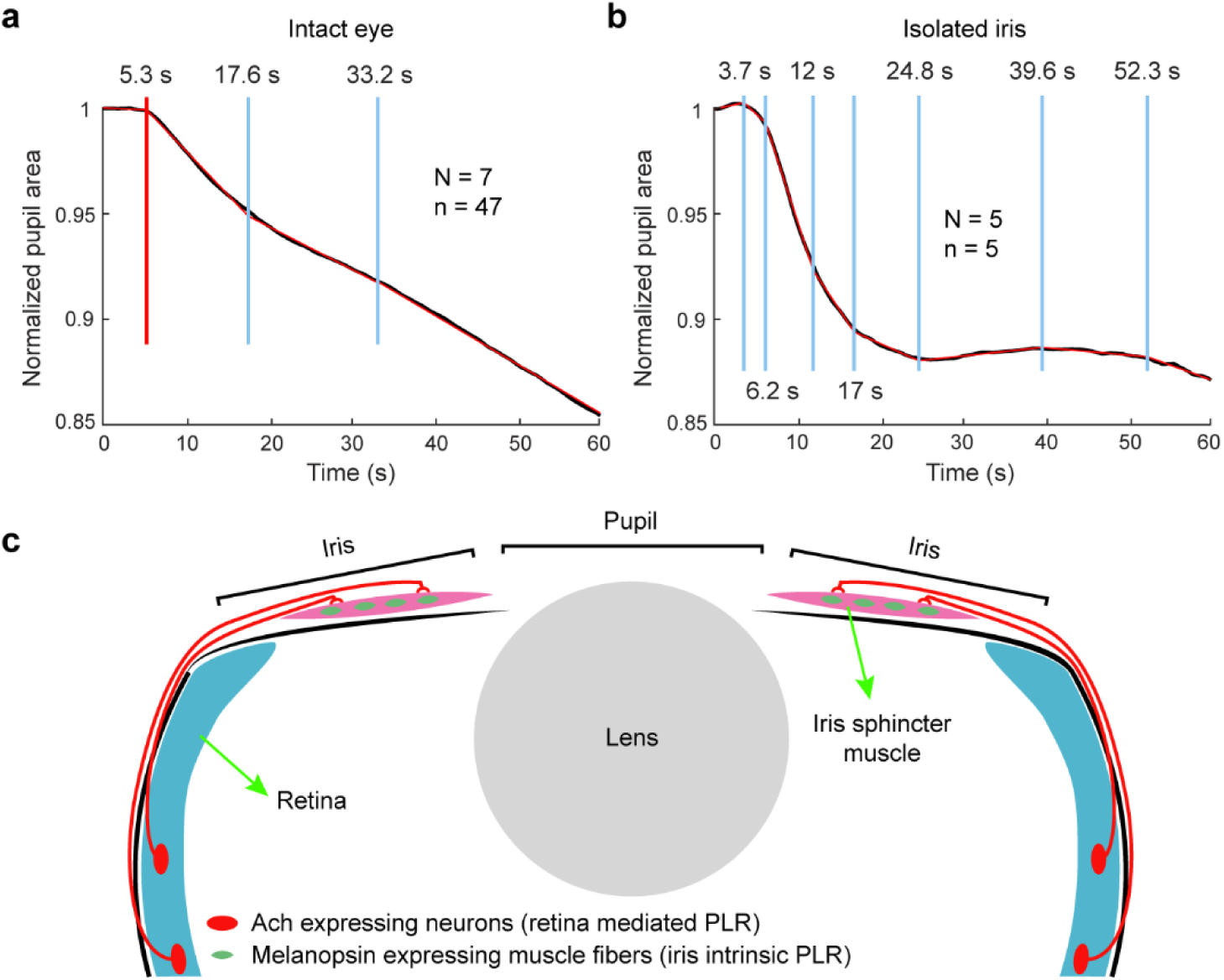
Temporal dynamics of the lamprey PLR and summarizing schematic. **a**, Graph showing the normalized pupil area average data of 7 animals (47 experiments) through time in response to light stimulation to the intact eye (black line), indicating the significant slope change points with blue vertical lines. The red line shows the fitted data. The red vertical line indicates the onset of muscle contraction. **b**, Graph showing the normalized pupil area through time in response to light stimulation to the isolated iris, indicating the significant slope change points with blue vertical lines. The black line shows the average data of 5 animals, whereas the red line shows the fitted data. **c**, Summarizing schematic. Muscle fibers in the iris sphincter muscle express melanopsin evoking an iris-intrinsic PLR. Additionally, cholinergic neurons project from the retina to the iris, mediating a second eye-intrinsic PLR mechanism. These cells potentially express melanopsin and/or are electrically coupled to melanopsin expressing cells.

## Discussion

Our results show that the PLR in lampreys is mediated by two mechanisms intrinsic to the eye, one iris-intrinsic mechanism mediated by melanopsin, and the other mediated by cholinergic projections from the retina (Fig. 5c). However, a brain-controlled PLR is absent. The lack of a brain-mediated mechanism explains the absence of a consensual PLR (Fig. 2h), which is a consequence of the bilateral innervation of the EWN from pretectum^1^. Additionally, the temporal dynamics of the PLR melanopsin component in other vertebrates explain the slow nature of the lamprey PLR^34^. In other vertebrates, the iris-mediated mechanism allows a sustained pupil contraction under strong steady light, which is not possible to achieve via the brain-mediated mechanism due to the small amount of light that reaches the retina through the pupil. In this manner, the highly photosensitive retinas of nocturnal/crepuscular animals are protected^4^. At least two types of photoreceptors, short and long, are present in the lamprey retina, and their response properties to light indicate that they are homologous to rods and cones, respectively^30,35–36^. Lampreys can live in deep waters, suggesting that they also have highly photosensitive eyes^37^. However, our results show that the iris-intrinsic mechanism alone is not sufficient to evoke a reliable PLR in these animals. Thus, a second mechanism, also mediated by melanopsin, evokes the PLR via cholinergic projections from the retina that, synergistically with the iris intrinsic mechanism, allows a consistent PLR. This is surprising given that both the iris-intrinsic and the retina-controlled mechanisms are mediated by melanopsin. However, it has been shown in mice that, although retinal and iridal melanopsin share a common phospholipase C-mediated phototransduction pathway, the downstream mechanisms are different^4^. It is likely that a similar situation is present in lampreys so that the iris and the retina-mediated mechanisms have different phototransduction dynamics, despite their common melanopsin activation, and therefore they complement each other allowing a robust and continuous pupil contraction.

The lack of a brain mediated mechanism is surprising given the high degree of conservation of the visual areas in the brain. The pretectum, which mediates the PLR in other vertebrates, shows conserved features from lampreys to mammals, including the same role mediating the optokinetic reflex^8^. Our results suggest that pretectal-mediated gaze stabilizing responses appeared earlier in vertebrate evolution than the pretectal-mediated PLR, and that the protection of the retina from light was initially based on eye-intrinsic mechanisms. Although the retina-mediated mechanism has been suggested in mice^2,7–9^, its contribution to the PLR is still controversial^5^ and, to our knowledge, our study is the first that demonstrates a direct projection from the retina to the iris. The lack of data in other vertebrates hinders a direct homology between the lamprey mechanism described here and the one proposed in mice. However, our results suggest that an iris-intrinsic PLR and one mediated by direct projections from the retina were present in early vertebrates and were conserved in some vertebrate groups, whereas the brain-mediated PLR appeared later in evolution.

## Methods

### Animals

Experiments were performed in 59 lampreys: 25 adult and 14 postmetamorphic sea lampreys (*Petromyzon marinus*), and 20 adult river lampreys (*Lampetra fluviatilis*). Animals were kept in enriched tanks with continuous oxygenation and filtration. All procedures were approved by the *Xunta de Galicia* under the supervision of the University of Vigo Committee for Animal use in Laboratory in accordance with the directive 2010/63/EU of the European Parliament and the RD 53/2013 Spanish regulation on the protection of animals use for scientific purposes. Minimizing the suffering and reducing the number of lampreys employed while maximizing the obtained data was a priority when designing the experiments.

### Experimental preparations

To investigate the brain contribution to the PLR, an isolated preparation of the brain with the eyes was used, so that electrophysiological experiments could be combined with pupil tracking and acute lesions or stimulation. For this, animals were deeply anesthetized prior to all dissections with tricaine methane sulfonate (MS-222; 100 mg/L; Sigma-Aldrich). Decapitation was performed between the third and fourth gills and the preparation was immediately immersed in a chamber containing refrigerated artificial cerebrospinal fluid (aCSF) with the following composition (in mM): 125 NaCl, 2.5 KCl, 2 CaCl_2_, 1 MgCl_2_, 10 glucose, and 25 NaHCO_3_, saturated with 95% O_2_/5% CO_2_ (vol/vol). All the muscles, viscera and skin were removed, exposing the eyes and the brain. Finally, the preparation was placed in a transparent chamber perfused with refrigerated aCSF. In this preparation, lesions were performed to analyze the encephalic contribution to the PLR. To analyze the presence of a putative Edinger-Westphal nucleus (EWN) the oculomotor nerves were cut and, to discard any possible encephalic contributions, the brain was totally removed. To investigate the presence of an eye/iris-intrinsic PLR, the eyes were dissected out from the preparation described above and placed in a transparent chamber perfused with ice-cooled aCSF. The iris was isolated by pinning down the eyes and cutting it out using small Castroviejo scissors. Then, the isolated iris was pinned down in a transparent chamber perfused with cold aCSF.

### Pupil tracking

The videos were recorded using digital camera USB-N&B NIR (IDS) coupled to a vari-focal manual iris lens with infrared detector, model T3Z2910CS-IR (Computar, CBC Group). All the videos were recorded in darkness, with the only constant source of light being an IR illuminator, model CM-IR56 (CMVision). To evoke pupil contraction, a white LED was used to apply light stimulation presented to one eye. First, the eye-brain preparation, isolated eye or iris was left in darkness for 20 min, and then light stimulation was applied during 5 min. These parameters were chosen to minimize the duration of the experiment while ensuring reliable pupil contraction and the viability of the preparation.

Iris contraction was tracked using DeepLabCut,^20^ a Python-based open-source software package to estimate poses through artificial neural networks. To achieve this, 20 key frames of each recorded video were selected randomly and four labels were placed at the external edge of the iris in each frame (Top, Right, Bottom and Left). Then, the neural network was trained and evaluated. When poor performance of the trained network was observed, based on its evaluation and/or visual inspection of the analyzed labeled videos, refinements were performed, and the network was further retrained. Subsequently, the trained network was used to extract the position of the labels throughout the videos.

### Electrophysiological recordings

Tungsten microelectrodes (∼1–5 MΩ) connected to a differential AC amplifier (model 1700, A-M systems) were employed to perform extracellular recordings of muscle and/or neuronal activity. Signals were digitized at 20 kHz using pClamp 10.4 software. Electrical stimulation was performed using borosilicate micropipettes (od = 1.5 mm, id = 1.17 mm; Hilgenberg) filled with aCSF connected to a stimulus isolation unit (MI401, Zoological Institute, University of Cologne). Microelectrodes and micropipettes were placed in the areas of interest employing micromanipulators (model M-3333, Narishige). To test the involvement of a putative EWN, electric stimulation of the oculomotor nerve was performed while video-recording pupil contraction and/or recording EMG activity in the iris. EMG activity in the dorsal rectus was used as a control (Extended Data Fig. 1a). In some cases, an incision was done in the iris to ensure that the recording electrode was in contact with the sphincter muscle. The stimulation intensity was tested from 0.01 to 1 mA, and both long stimulation trains with short pulses (30 s stimulation, 10 ms duration pulses, 10 Hz) and single pulse stimulations with long duration (10 s) were tested. To ensure that the lack of responses was not due to invalidity of the preparation, extracellular recordings in reticulospinal neurons in response to optic tract stimulation were carried out as control.

### Drug applications

The role of melanopsin mediating the iris-intrinsic PLR was tested using AA92593 (60 µM; MedChemExpress), a selective and reversible antagonist of melanopsin-mediated phototransduction without affecting rod- and cone-mediated responses. AA92593 was bath-applied to the iris, isolated as previously described. To assess cholinergic innervation to the iris, acetylcholine (ACh) 100 µM (Sigma-Aldrich) was bath applied to the isolated iris, while monitoring pupil contraction. A fragment of extraocular muscle was also used as a control to test ACh activation. Atropine (1 mM; Sigma-Aldrich) was employed to block cholinergic transmission both bath-applied to the isolated iris, and to isolated eyes. In the latter case, bath-applied administration was combined with atropine injections in the eye to ensure the exposure of the iris muscle to the drug. To analyze the existence of gap-junction between iris muscle fibers, octanol (500 µM; Carlo Erba) was used as a reversible blocker.^5^ Experiments were performed in the isolated iris. In all cases, drug effects were reversed (partially or totally) washing out with clean aCSF.

### Anatomical tract tracing

Neurobiotin (Vector Laboratories) or dextran-rhodamine (tetramethylrhodamine, 3,000 Da, Invitrogen, D3308) was used to study the neuronal circuits involved in the PLR. Tracer crystals were applied in the cut oculomotor nerve, and/or the iris. For tracer application in the iris, either lesions with a sharp needle or an opening using Castroviejo scissors was made to ensure the direct application to the muscle. After the tracer applications, the samples were thoroughly washed and submerged in aCSF at 4°C for 1 to 4 days in darkness to allow the transport of the tracer. Then, the preparations were fixed in 4% formaldehyde and 14% saturated picric acid in 0.1 M phosphate buffer (PBS), pH 7.4, for 12-24 h, and cryoprotected in 20% (wt/vol) sucrose in PBS for 4-12 h. Afterwards, they were embedded in OCT compound (Tissue-Tek, Sakura), cut in transverse sections of 30 μm thickness in a cryostat (Leica cm1950) and collected on gelatinized slides. To detect neurobiotin, sections were incubated with Cy2-conjugated streptavidin (Alexa Fluor 488, 10,000 Da, Invitrogen, D22910) 1:1000 in blocking solution (1% bovine serum albumin, 0.1% sodium azide and 0.3% Triton X-100 in PBS). The sections were subsequently mounted in DAPI-containing glycerol (DAPI Fluoromount-G, SouthernBiotech). For some injections, neurobiotin was detected using the ABC kit (SIGMA), and the labeling was observed flat-mounting fragments of the retina. For this, retina fragments were fixed in 1% glutaraldehyde, 4% paraformaldehyde and 15% saturated picric acid in PB for 24 h. Samples were placed over a slide and covered with melted agar 4%. Once agar solidified, the samples were immersed again in melted agar a 4% and horizontally sectioned in a vibratome. 100-200 µm thick sections were obtained and subsequently incubated in 1% sodium borohydride in PBS for 1 h, in 15% H_2_O_2_ in PBS for 1h, and 1% bovine serum albumin, 0.3% Triton X-100 in PBS for 1 h. Sections where then incubated in ABC kit, diluted 1/100 in PBS for 1 h, washed in PBS three times and developed with 0.5% of 3,3’-diaminobenzidine (SIGMA) and 0.01% H_2_O_2_ in PB for about 15 min. Sections were mounted with glycerol for observation.

### Hematoxylin-eosin staining

Eyes were fixed in buffered 4% paraformaldehyde and 15% of saturated picric acid, and paraffin embedded following a standard protocol. 8 µm thick sections were deparaffined, hydrated, and stained with Mayer hematoxylin and eosin Y (0.2%). After the staining, sections were dehydrated, cleared and coverslipped with Eukitt (ORSAtec). Hematoxylin-eosin staining was also performed in cryostat sections, obtained as described above.

### Electron microscopy

Adult and postmetamorphic lamprey eyes were dissected out and fixed in 1% glutaraldehyde, 4% paraformaldehyde and 15% saturated picric acid in PB for 24 h. They were osmified in 1% of osmium tetroxide in PB for 30 min, dehydrated in ethanol and embedded in Durcupam (Fluka). Ultrathin sections were obtained in an ultramicrotome (Reitcher), placed onto copper grids, and contrasted with uranyl acetate and lead citrate. Before cutting ultrathin sections, some semithin sections (0.5 µm) were obtained, placed onto slices, stained with toluidin blue, and observed and photographed at light microscopy.

### Immunofluorescence

Eyes from postmetamorphic and adult lampreys were fixed in 4% paraformaldehyde in PB for 24 hours. After that, the caudal half of the ocular globe and the eye lens were removed, trying to keep the lens covering tissue. Some meridian cuts were done in the remaining of the ocular globe to get extended and nearly flat samples. These samples were then incubated in 1% sodium borohydride in PBS for 1 h, and 1% bovine serum albumin, 0.3% Triton X-100 in PBS for 1 h. After that, they were immersed in a monoclonal anti-mouse acetylated tubulin (SIGMA) diluted 1/1000 in PBS for 16-18 h. Samples were washed several times in PBS and incubated in a goat anti-mouse conjugated with Alexa fluor 488 (Invitrogen) diluted 1/100 in PBS for 1 h. After several washes in PBS, samples were placed and extended onto slides, coverslipped with anti-Fade reagent (Molecular probes), and observed and photographed at fluorescence microscopy.

### Image analysis

Photomicrographs were taken with a digital camera (Nikon DS-Ri2) coupled to a Nikon ECLIPSE Ni-E fluorescence microscope. Confocal images were obtained with a Leica Stellaris 8 microscope 510. Images were processed using ImageJ 1.53k and GIMP 2.1. Images were only adjusted for brightness and contrast. Figures were made using Adobe Illustrator CC 2019.

### Quantification

Throughout this paper the number of animals (N) and the number of experiments performed (n) are indicated where applicable. Data analysis was performed using custom written functions in Matlab R2020b. To calculate pupil contraction, videos were first analyzed to extract the positions of the four labels above described using DeepLabCut. Then, the distances between the labels in the top and bottom edges of the pupil, and the right and left edges were used to calculate the radius in the X and Y axes of the pupil, respectively. This was in turn used to calculate the area of the pupil applying the formula for an ellipse. Area data were normalized to the first frame of light stimulation or drug application (for ACh application) to calculate the percentage of reduction. To detect significant slope changes, we used the Matlab function *findchangepts*, which returns the index at which the mean of x changes most significantly.

### Statistical analysis

Statistical analysis was done using JASP 0.16. Paired t-tests were used to compare pupil reduction between different conditions. Throughout the Figs., sample statistics are expressed as means ± SD. The degree of statistical significance is indicated as follows: *P < 0.05, **P < 0.01, ***P < 0.001.

## Acknowledgements

We are grateful to Sten Grillner and Manuel A. Pombal for their constant support and valuable comments on the manuscript, to Daichi G. Suzuki and Brita Robertson for his valuable comments on the manuscript, to Martiño Barreiro and Emma Rodríguez for technical support, to Eduardo Pena for providing hardware, and to Tobias Wibble and the river station of A Freixa for helping with lamprey supply. This work was supported by Proyectos I+D+i PID2020-113646GA-I00 funded by MCIN/AEI/ 10.13039/501100011033 and by “ERDF A way of making Europe”, the Ramón y Cajal grant RYC2018-024053-I funded by MCIN/AEI/ 10.13039/501100011033 and by “ESF Investing in your Future”, Xunta de Galicia (ED431B 2021/04 to JPF and ED481A 2022/433 to CNG), and CINBIO.

## Author contributions

Conceptualization: CJL, JPF; Experimental design: CJL, PRR, MM, JPF; Data acquisition: CJL, PRR, MB, CNG, MM; Data analysis: CJL, PRR, MB, CNG, MM, JPF; Writing: CJL, MM and JPF with inputs from all the authors; Supervision and Funding acquisition: JPF.

## Extended Data

**Extended Data Fig 1.**
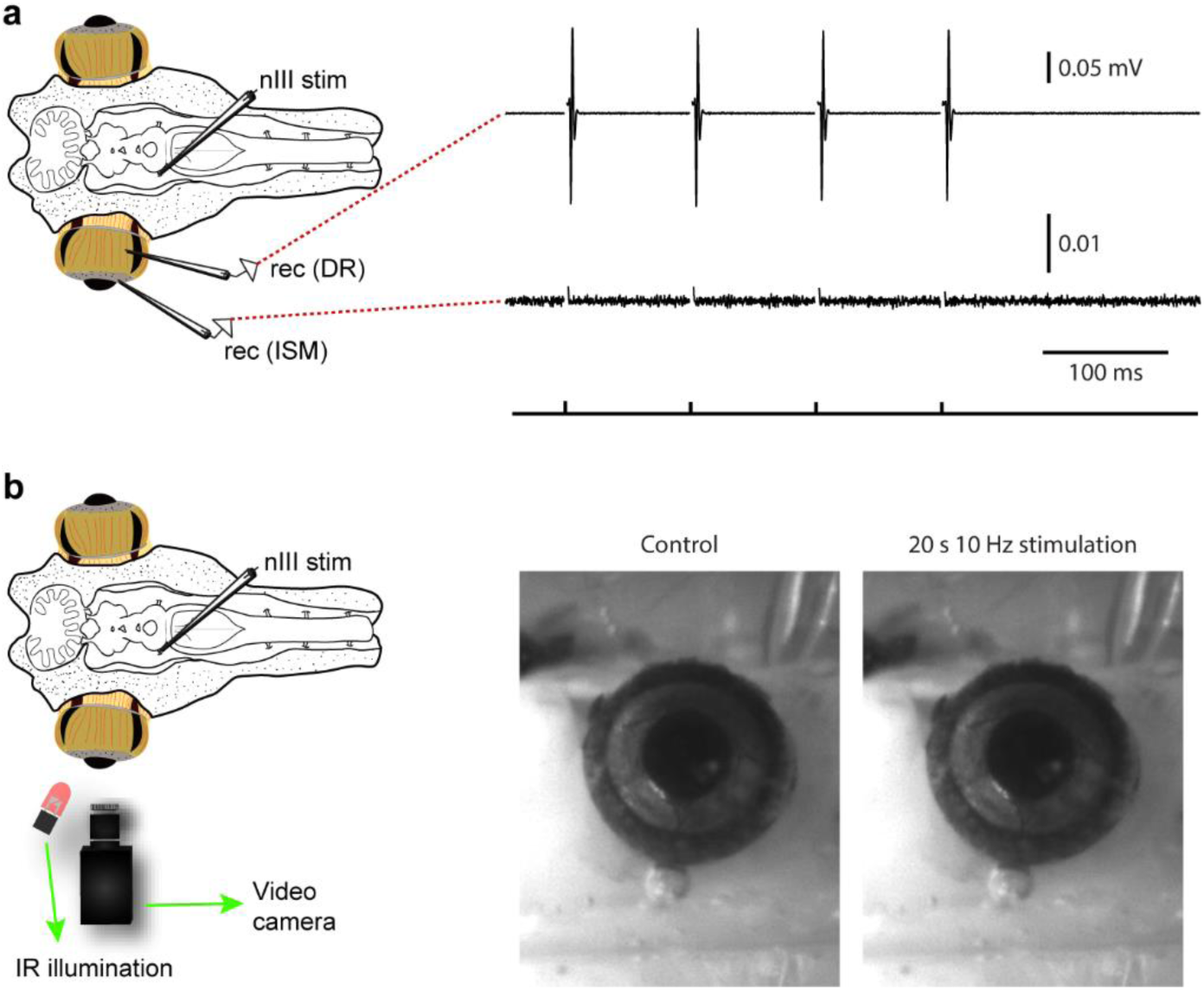
Electric stimulation of the nIII does not evoke PLR in lampreys, related to Fig. 2. **a**, EMG activity recorded in the dorsal rectus (DR, top) as a control, whereas no responses are observed in the iris sphincter muscle (ISM, bottom) after a four pulses electric stimulation of the oculomotor nerve (nIII; 10 Hz). **b**, Representative frames showing no changes in pupil size before (left, control), and after (right) electric stimulation of the nIII (20 s, 10 Hz).

**Extended Data Fig 2.**
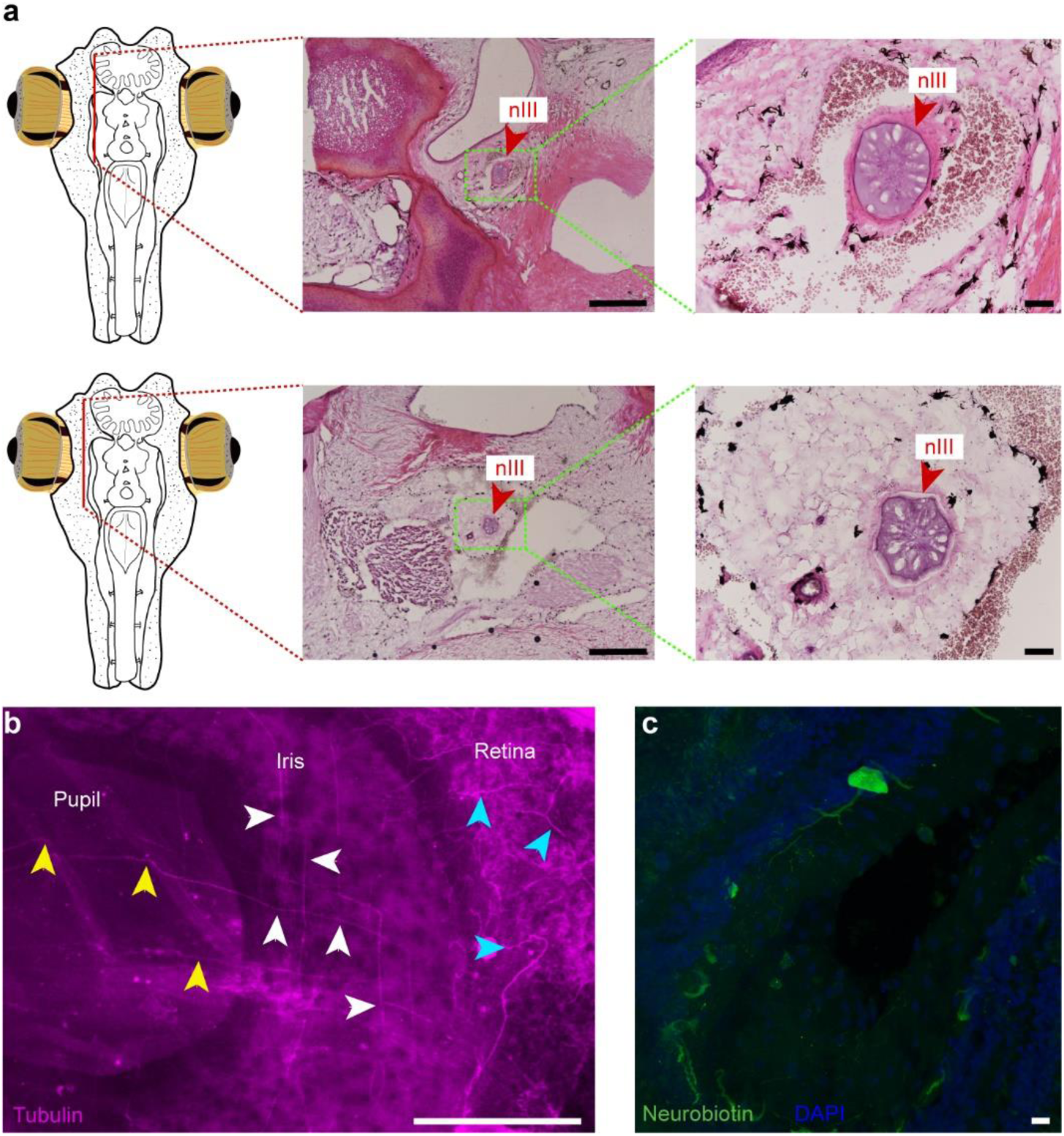
Iris axonal innervation originates in the retina, related to Fig. 4. **a**, Representative hematoxilin-eosin stained sagittal sections of the head, following transversally the oculomotor nerve (nIII) from its exit in the brain. The top sections show the nerve close to the brain, at its crossing level of the cartilage surrounding the brain. The bottom images show the nerve closer to the eye. **b**, Photomicrograph showing the expression of acetylated tubulin in a flat-mounted eye. Labeled axons can be seen coursing from the retina to the iris. Axons are indicated with blue arrowheads at the level of the retina, white arrowheads at the level of the iris, and yellow arrowheads at the level of the pupil. **c**, Retrogradely labeled cell found in the inner nuclear layer (INL) after neurobiotin injection in the iris. Scale bars = 1 mm in **a** (left); 100 µm in **a** (right); 500 µm in **b**; 10 µm in **c**.

